# CyDAP–A fluorescent probe for cytosolic dopamine detection

**DOI:** 10.1101/2020.09.04.283911

**Authors:** Jing-Yi Jeng, Lee Sun, Jia-Chi Wang, Cheng-Yuan Lin, Chih-Ping Hung, Li-An Chu, Hui-Yun Chang, Ann-Shyn Chiang, Tzu-Kang Sang

**Author notes:** equal contribution (Jing-Yi Jeng, Lee Sun). Correspondence: Tzu-Kang Sang, Ph.D., Institute of Biotechnology and Department of Life Science, National Tsing Hua University, 101, Section 2, Kuang-Fu Road Hsinchu 30013, Taiwan.

## Abstract

Dopamine (DA) is an essential neurotransmitter modulating motor and cognitive functions. Several neurological disorders, including Parkinson’s disease (PD) and drug addiction, are the result of DA system dysfunction; however, it remains incomplete understood of why DA neuron is selectively more vulnerable than other neurons. Here we utilize the spectral feature of human MAO B (monoamine oxidase B) to design a genetic-amenable, GFP-based fluorescent probe CyDAP. Upon genetic and pharmacological manipulations to elevate the cytosolic DA levels in cells and *Drosophila* models, CyDAP shows enhanced GFP emission, suggesting this probe is feasible for DA detection. Furthermore, we observe that expressing human α-Synuclein in *Drosophila* elicited GFP emission from CyDAP, suggesting a link between cytosolic DA imbalance and regional vulnerability in PD context. Importantly, CyDAP can detect the change of cytosolic DA in live *Drosophila* brains, as demonstrated by time-lapse and the 4D light-sheet confocal recording. CyDAP may serve as a tool for evaluating metabolic deregulation of DA in brain models of PD and other DA system-related psychiatric disorders.

## Introduction

Neurotransmitter dopamine (DA) modulates motor and cognitive functions, and the impairment of this system causes a wide range of neurological conditions, including Parkinson’s disease (PD), psychiatric disorders, and drug addiction (Liss and Roeper, 2008). The precursor of DA is L-DOPA (L-3-4-dihydroxyphenylalanine), which is converted from L-tyrosine by TH (tyrosine hydroxylase) and its cofactor. The subsequent L-DOPA decarboxylation, catalyzed by AADC (aromatic amino acid decarboxylase, also known as dopa decarboxylase, or DDC) yields DA (Carlsson et al., 1958; Christenson et al., 1972; Holtz P, 1938; Nagatsu et al., 1964). The newly made neurotransmitters are transferred into the vesicle by the vesicular monoamine transporter (VMAT) (Liu et al., 1992) to be released upon the fusion of the synaptic vesicle and the cell membrane. The released DA could be recycled through DA transporter (DAT) (Giros et al., 1996), thereby reload the neurotransmitter back into the vesicles. Alternatively, the recycled DA could be catalyzed by monoamine oxidase (MAO) and catechol-O-methyltransferase (Eisenhofer et al., 2004; Hare, 1928; Westlund et al., 1985). Because of the unstable nature of the catechol ring, free cytosolic DA is subject to oxidation, which could produce the toxic DA quinone and other catabolic adducts (Fornstedt et al., 1990; Sulzer and Zecca, 2000).

The degeneration of nigrostriatal DA neurons is the pathological hallmark of PD of which impairment vindicates the primary motor symptoms, such as tremor and slow movement in the patients (Przedborski, 2017). While various genetic and environmental factors risk PD pathogenesis, little is known why DA neurons are more vulnerable than others (Surmeier et al., 2017). A plausible scenario proposed is DA itself, which ignites selective vulnerability (Conway et al., 2001; LaVoie et al., 2005). However, experimental support of this suggestion has been limited because monitoring DA dynamics with the subcellular resolution is challenging. The common ways to analyze DA is either by high-performance liquid chromatography (HPLC), which measures DA from tissue homogenates (Hjemdahl, 1984), or by fast-scan cyclic voltammetry, which inserts a carbon fiber electrode (CFE) to read the electrical outputs upon DA chemical reaction (Robinson et al., 2003). However, both approaches quantify DA from tissues as a whole. A modified electrochemical technique applied CFE for intracellular patch recording to report the cytosolic pool of catecholamine in cultured cells and brain slices (Mosharov et al., 2003; Mosharov et al., 2009). Utilization of fluorescent false neurotransmitters could visualize the mimicked DA release from the synapse (Gubernator et al., 2009), yet whether it could be used for analyzing cytosolic DA dynamics is unclear.

MAOs catalyze the oxidation of biogenic and dietary amines like DA (Bach et al., 1988; Binda et al., 2002; Miller et al., 2000). The human genome contains two distinct MAO genes, A and B, which encode proteins that share a ~73% sequence identity with a similar structure (Bach et al., 1988; Edmondson et al., 2004). Both enzymes function as homodimers and anchor onto the mitochondrial outer membrane by their C-terminal helices (Binda et al., 2006; Mitoma and Ito, 1992). Despite the high similarity, the geometry and size of their substrate cavities are different (Edmondson et al., 2007), which account for the preferential affinity. For example, MAO A catalyzes serotonin more effectively than MAO B, whereas MAO B performs better than MAO A in catalyzing DA (Edmondson et al., 2009). These flavoproteins require flavin adenine dinucleotide (FAD) as a cofactor to process substrates. Importantly, MAO B has a unique characteristic spectrum (Li et al., 2006). With its cofactor in an oxidized state, MAO B absorbs blue light (400-500 nm). This spectral property is receded when FAD is reduced to FADH2 upon substrate binding. Therefore, we envisage that the oxidized form of MAO B might absorb GFP emission (~500 nm) but GFP fluorescence can be emitted when MAO B is reduced after substrate binding, i.e., when there is bounded DA. Such shifting may serve as an optical means to report the presence of DA in the subcellular domain.

Currently, the available method for detecting cytosolic DA is limited to the electrochemical technique requiring an invasive approach. Here, we show a genetic amenable fluorescent probe made by MAO B and GFP chimera that is capable of detecting cytosolic free DA *in vivo.* We demonstrate the GFP signals emitted from the probe are closely associated with cytosolic DA levels through the genetic and chemical manipulations, and the probe is feasible for detecting DA in living brains. By incorporating this probe in the *Drosophila* model expressing α-Synuclein or its disease-associated allele, we show a significant increase of GFP emission, suggesting this notorious PD gene may alter cytosolic free DA pool. This probe could be a useful tool to decipher the puzzle of regional vulnerability in PD and DA-related disorders.

## Results

### A DA Probe Uses the Light Absorption Feature of Human MAO B

We initially tested three constructs (MG-s, MG-m, and MG-l) by fusing AcGFP (*Aequorea coerulescens* GFP: ex. 475 nm/ em. 505 nm) to the C-terminal of truncated MAO B (Figure S1A and S1B). Structural simulation of these MG probes estimated that the distance between the chromophore residue of GFP (Tyr62 of AcGFP) and the flavin-binding site of MAO B (Cys397) is ~38 Å, within the range (<100 Å) of intramolecular fluorescence resonance energy transfer (FRET, Figure S1A). We tested the probes in *E. coli* and *rat pheochromocytoma* (PC12) cells. A Western analysis using the anti-GFP antibody confirmed all MG probes preserve the GFP epitope (Figure S1C). However, GFP fluorescence was undetectable in cells expressing either MG-s or MG-m and only weak signals shown in MG-l (Figure S1D), suggesting that MAO B might absorb the emitted GFP signals. We thus named this phenomenon the “shielding effect.”

To test if the shielding effect could be reversed upon substrate binding, we transfected MG-s into PC12 and examined the GFP signal change before and after DA treatment. To facilitate DA uptake, we followed a previous report by differentiating PC12 cells with nerve growth factor (NGF) (Mudumba et al., 2002) to induce DAT expression (Figure S1E). Flow cytometry analysis revealed approximately eleven folds increase of GFP-positive cells after NGF plus DA treatments compared with untreated control (Figure S1F), supporting that DA supplement could reverse the shielding effect of MG-s. To visualize this effect directly, we injected DA into MG-s-transfected PC12 cells and observed flux of GFP fluorescence (Supplementary Video 1-2 and Figure S2). Together, we demonstrated the unique light-absorption feature of human MAO B applies to engineer a fluorescent probe for DA detection.

### DA Probe with a Comparable MAO B Activity and Shielding Effect

While MG-s probe enabled DA detection, MAO B activity was low (Figure S3B). As this might affect the fidelity of DA detection, we modified the probe aiming to raise the enzyme activity while still retaining the shielding effect. Within the tested constructs (Figure S3A), MAO B activity assays showed GFP fused to the N-terminus of MAO B (GM5) preserved enzyme activity comparable to normal MAO B (Figure S3B), but lacked the shielding effect (Figure S3C). Constructs with GFP fused to the C-terminus of MAO B (M4G and M5G), on the other hand, encountered low enzyme activity, similar to MG-s (Figure S3B). Guided by structure simulation, we made constructs by replacing a short segment of MAO B C-terminal peptides with GFP (MGM1, MGM2, and MGM3). Despite retaining the mitochondrial anchorage domain, these constructs were essentially lack of MAO B activity (Figure S3B). Together, we postulated that the integrity of the C-terminus of MAO B is critical to the enzyme activity.

To ensure the minimal change of MAO B, we adopted the split-GFP system to modify the probe. By incorporating the 11th β-sheet of the super folder GFP (GFP11, aa. 215-230 of AFC90853 (Pedelacq et al., 2006)) to the C-terminal-end of wild-type MAO B, this construct, MsfG, preserved ~65% of MAO B activity (Figure S3B). We then co-expressed MsfG and GFP1-10 (aa. 1-214 of super folder GFP, hereafter refer to sfG110) in cells to test the shielding effect. Although GFP signals were not detected (Figure S3C), we were concerned that this was because split-GFP fails to reconstitute. It is possible that the C-terminal hydrophobic helix of MAO B ends in the mitochondrial intermembrane space as speculated previously (Binda et al., 2004) and thus cannot form full-length GFP with cytosolic sfG110. If the C-terminal GFP11 from MsfG fails to expose the cytosolic side of the mitochondrial outer membrane, it could hinder its reconstitution with cytosolic sfG110 and generating false-positive shielding effect. Therefore, we added the second transmembrane domain of human Mitofusin 2 (Mfn2; aa. 627-648) to the C-terminal end of MAO B, before sticking GFP11 fragment. This modified construct, named MMG1, was predicted to have two hydrophobic domains (Figure 1A). The structural simulation showed the estimated distance between the Tyr residue of GFP11 (RDHMVLHE<u>Y</u>VNAAGIT) and the flavin-binding site of MAO B (MAO B^C397^) is ~17.68 Å (Figure 1B), within the permitted distance for FRET to occur.

**Figure 1.**
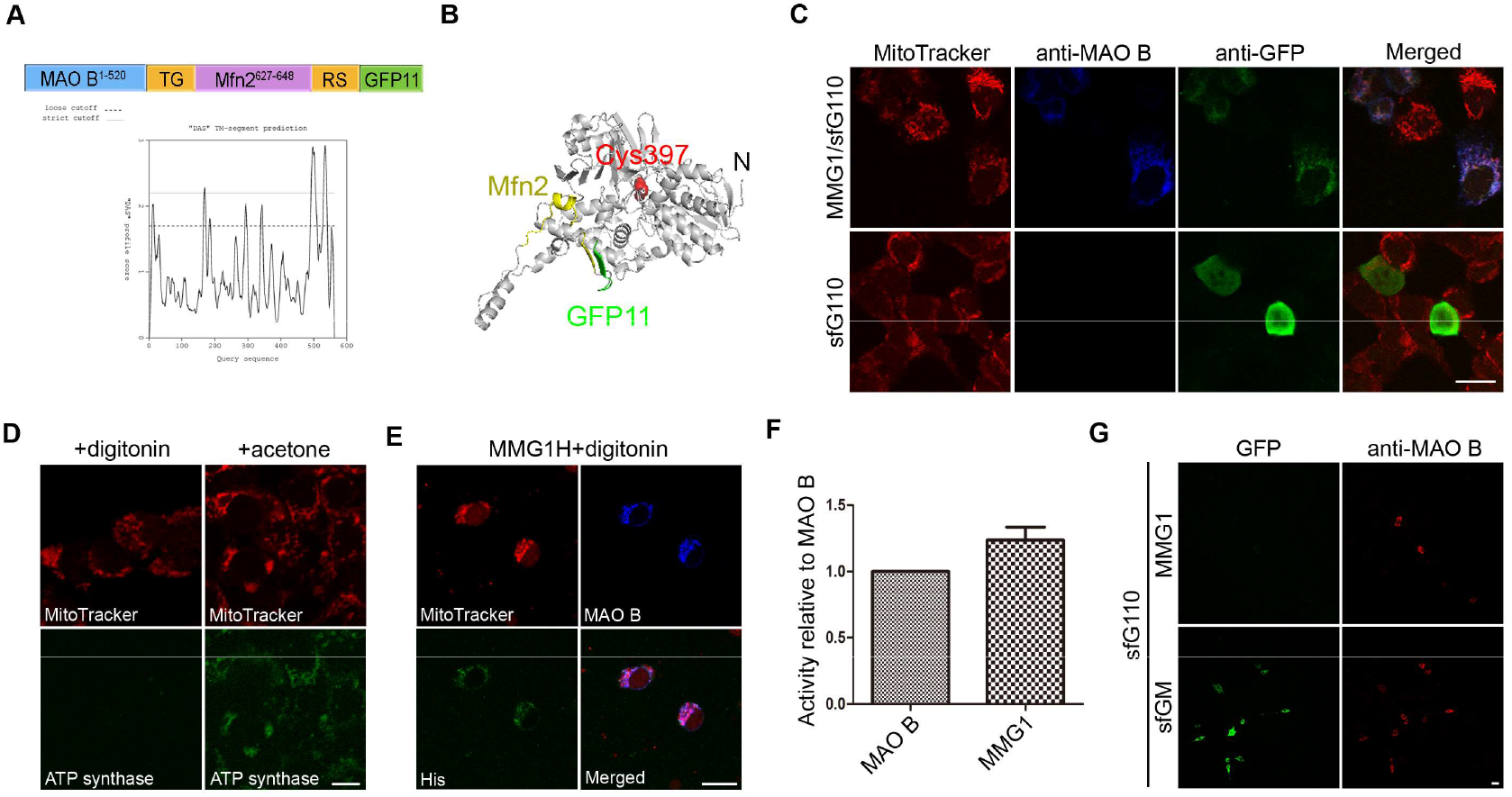
Characterization of DA probe MMG1. (A) Schematic depiction of MMG1 with the corresponded fusion components and linker sequences (upper panel). The prediction of MMG1 transmembrane segments, a.a. 490-509 and a.a. 527-540 (lower panel; http://www.sbc.su.se/~miklos/DAS/), are corresponded to the C-terminal peaks of the linear protein sequence. (B) The prediction of MMG1 structure by I-TASSER (https://zhanglab.ccmb.med.umich.edu/I-TASSER/). PyMOL highlights the Mfn2 (yellow) and GFP11 (green) fragments within MMG1. A red ball-and-stick molecule marks the FAD-binding residue Cys^397^ of MAO B. (C) Confocal images of HEK293 cells transfected with the indicated constructs stained with MitoTracker (red), anti-MAO B (blue), and anti-GFP (green). Notice the GFP signals in the upper panels are sequestered to the mitochondria. (D, E) Confocal images of HEK293 cells transfected with His-tagged MMG1 (E) and permeabilized with the indicated reagents. Cells are stained with MitoTracker (red), anti-ATP synthase (green in D), anti-MAO B (blue), and anti-His (green in E). (F) MAO B activity of the indicated probes. Data are normalized to MAO B control from three independent assays. (G) Confocal images of PC12 cells co-transfected with sfG110 and the indicated constructs. Cells are stained with anti-MAO B (red). GFP channel shows intrinsic signals. Scale bars: 10 μm.

To experimentally validate MMG1 topology, we co-expressed MMG1 and sfG110 in HEK293 cells. Immunostaining of cells expressing sfG110 alone showed ubiquitous anti-GFP signals in the cells; however, we found that anti-GFP signals preferentially labeled mitochondria (marked by MitoTracker), which were also stained by anti-MAO B (Figure 1C), suggesting the reconstitution of the split GFP of MMG1 and sfG110. To confirm the GFP reconstitution occurred in the cytosol, we made a His-tagged MMG1 at its C-terminus (MMG1H) and selectively permeabilized the plasma membrane with 20 μM digitonin. Unlike acetone, this digitonin concentration does not permeabilize the mitochondrial membrane, thus preventing antibody detection of ATP synthase on the inner-mitochondrial membrane (Figure 1D). We were able to label MMG1H with an anti-His antibody of which signals were co-localized with MAO B in mitochondria (Figure 1E), supporting the C-terminal tail of MMG1 was exposed in the cytosol to enable GFP reconstitution. Importantly, MMG1 preserved a comparable enzyme activity like the native MAO B protein (Figure 1F). Comparing the isolated mitochondria from cells expressing either MMG1 or MAO B revealed similar affinity (Km=451.8 ± 83.88 μM for MMG1, and 369.5 ± 117.9 μM for MAO B) toward MAO B substrate benzylamine (BZA, (Sourkes, 1980)), indicating MMG1 functions like MAO B.

Next, we tested the shielding effect of MMG1. We generated a split version of GM5 (Figure S3C) by attaching GFP11 to the N-terminus of MAO B (sfGM) to serve as a control for GFP fluorescence. Cells co-expressing sfGM and sfG110 showed robust GFP signals, similar to GM5. On the contrary, GFP signals were virtually undetectable in cells co-expressing MMG1 and sfG110 (Figure 1G). Together, these results demonstrated that MMG1 fits the required criteria preserving a comparable enzyme activity of MAO B and the shielding effect.

### MMG1/sfG110 Detects DA in Culture Cells

To test the probe for DA detection, we co-transfected the corresponded plasmids to PC12 cells. As expected, the shielding effect prohibited GFP emission in cells co-expressing MMG1/sfG110, but not to sfGM/sfG110 control. Upon treating L-DOPA to cells transfected by MMG1/sfG110, we observed GFP signals in a dose-dependent manner, suggesting the reverse of the shielding effect (Figure 2A and 2B). Quantitative analysis by flow cytometry confirmed that more cells were emitting GFP with higher L-DOPA levels (Figure 2C). To validate the reversion of the shielding effect is FAD-dependent, we made a FAD-binding mutant MMG1^C397A^ (Rebrin et al., 2001). Using a DAT-expressing HEK293T cell line, we compared the GFP emission of cells expressing either MMG1/sfG110 or MMG1^C397A^/sfG110 after treating with MAO B substrate BZA. Consistent with DA-treatment, cells expressing MMG1/sfG110 showed GFP emission, whereas MMG1^C397A^/sfG110 expressing cells did not (Figure 2D). Together, these data support MMG1/sfG110 probe is feasible for DA detection in a FAD-dependent manner.

**Figure 2.**
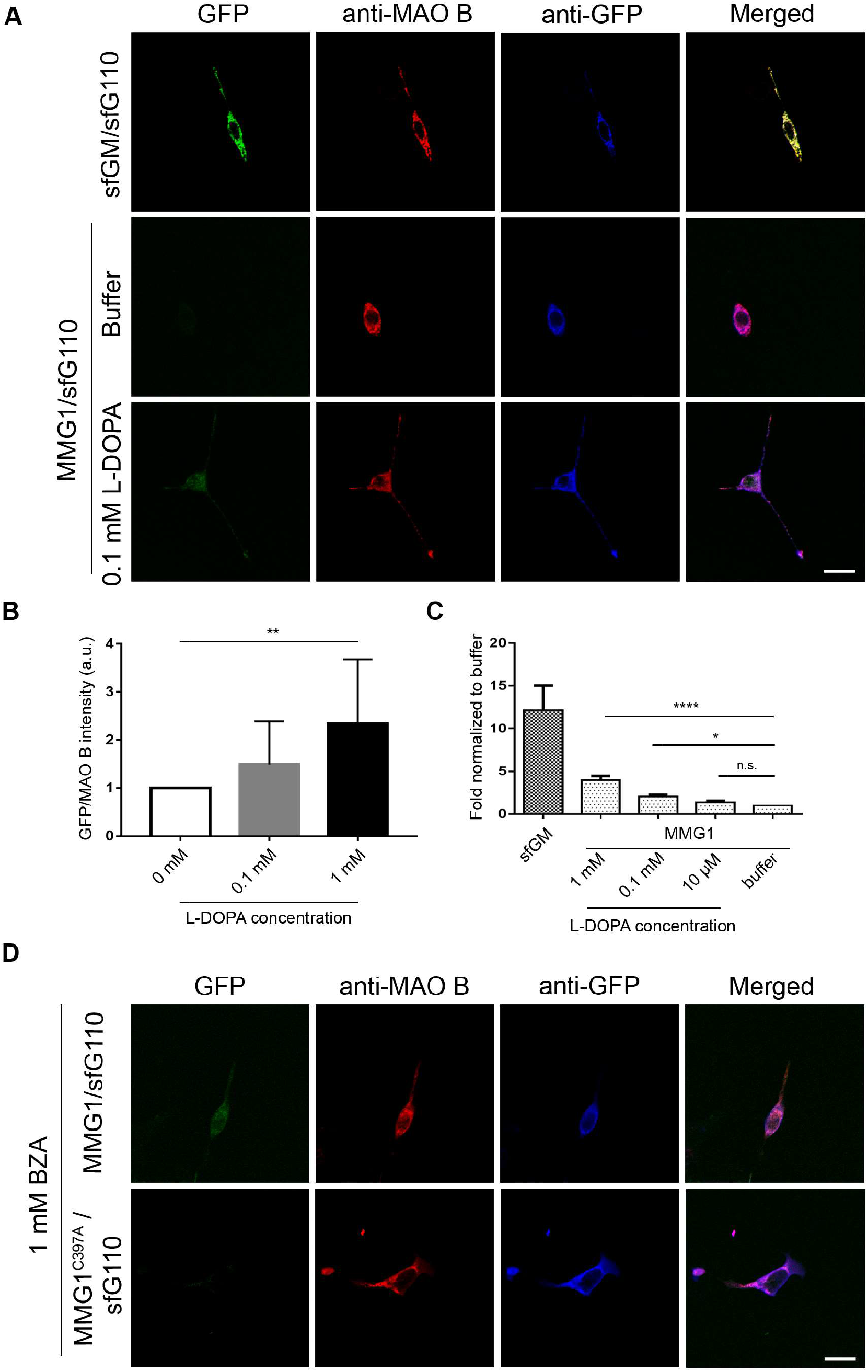
MMG1/sfG110 reacts to MAO B substrates by emitting GFP fluorescence. (A) Confocal images of differentiated PC12 cells transfected with the indicated constructs stained with anti-MAO B (red) and anti-GFP (blue). GFP channels (green) are intrinsic signals. The lower panels show cells expressing the indicated constructs are treated with 0.1 mM L-DOPA. (B) Quantitation of GFP intensity from cells expressing MMG1/sfG110 treated with different concentrations of L-DOPA. GFP pixels are normalized to anti-MAO B-labeled pixel in cells, n≥12. (C) Flow cytometry analysis of GFP-positive cells co-expressing sfG110 and the indicated constructs. Cells are treated with different concentrations of L-DOPA, n≥4. Values shown represent mean ± SE. n.s., not significant; *, p < 0.05; **, p < 0.01; ****, p < 0.0001 as compared to non-treatment (0 for B and Buffer for C) within the MMG1-expressing cells (one-way ANOVA with Dunnett’s multiple comparisons test). (D) Confocal images of HEK293T cells co-transfected with sfG110 and the indicated MMG1 variants. Cells are treated with 1mM benzylamine (BZA) and stained with anti-MAO B (red) and anti-GFP (blue). GFP channels (green) are intrinsic signals. Notice cells expressing FAD-binding mutant MMG1^C397A^ do not emit intrinsic GFP signals. Scale bars: 10 μm.

### MMG1/sfG110 Detects Endogenous DA in the *Drosophila* Brain

To test the efficacy of MMG1/sfG110 in the brain, we used *Drosophila* as a model. DA neurons in *Drosophila* brain produce, store, and recycle DA similar to the mammal. However, unlike mammals, fly processes DA by acetylation and alanylation (Binda et al., 2003; Hintermann et al., 1996; Tracy L. Paxon, 2005; Yamamoto and Seto, 2014). Fly genome lacks MAO orthologous; therefore, the endogenous interference by this enzyme is neglectable.

We harnessed the bipartite GAL4/UAS system in which the generated UAS transgenic flies were bearing MMG1 and related transgenes, which could produce the encoded proteins upon introducing the cell type-specific GAL4 driver. Using *TH-GAL4* to express either MMG1/sfG110 (*TH>MMG1/sfG110)* or sfGM/sfG110 in DA neurons, we confirmed their localization to the mitochondria where immunostaining of anti-MAO B and anti-ATP5a (a mitochondrial complex V protein) signals were overlapped (Figure 3A). Consistent with the culture cell result, DA neurons expressing sfGM/sfG110 showed robust GFP signals, whereas MMG1/sfG110 cells only emitted weak GFP fluorescence (Figure 3A).

**Figure 3.**
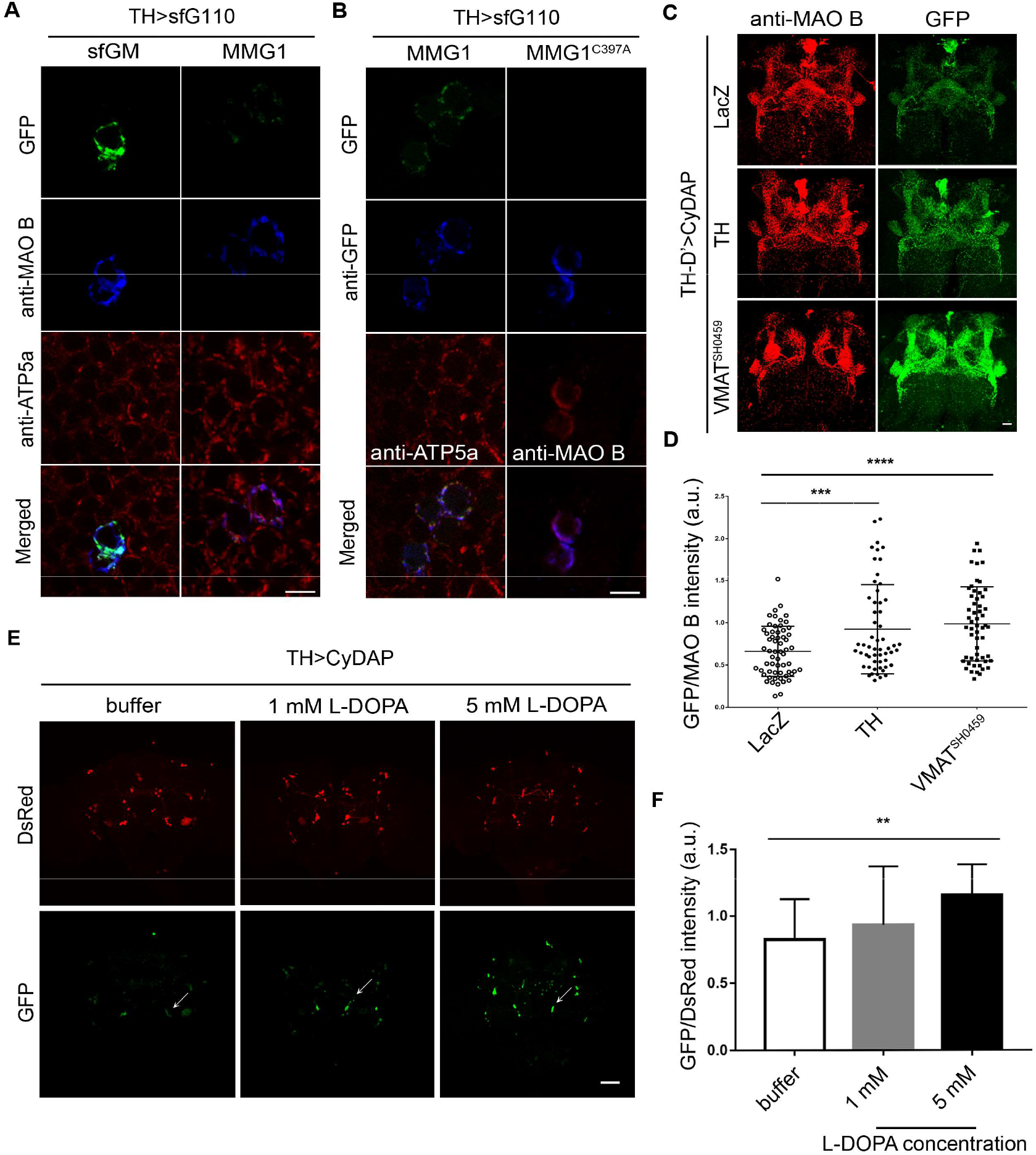
CyDAP responses to DA dynamics in *Drosophila* brains. (A, B) Confocal images of *TH>sfG110* flies co-expressing the indicated constructs. Cells are stained with anti-MAO B (blue in A, red in the right panel of B), anti-ATP5a (red in A and the left panel of B), and anti-GFP (blue in B). GFP channels (green) are intrinsic signals. Notice weak intrinsic GFP fluorescence in MMG1-expressing cells, but not in MMG1^C397A^ (B). (C) Confocal images of *TH-D’>CyDAP* flies co-expressing LacZ control (upper panels), TH (middle panels), or in VMAT heterozygous background (VMAT^SH0459/+^, lower panels) stained with anti-MAO B (red). GFP channels (green) are intrinsic signals. (D) Quantification of GFP intensity after normalized to anti-MAO B signals. Four neuropils innervated by TH-D’ neurons in each brain from a total of 14 flies of each group are measured. Values shown represent mean ± SE. ***, p < 0.001; ****, p < 0.0001 as compared to lacZ control (one-way ANOVA with Dunnett’s multiple comparisons test). (E) Confocal images of *TH>CyDAP* flies co-expressing DsRed fluorescent protein. Flies are fed with buffer used to dissolve L-DOPA or the indicated levels of L-DOPA. Notice that the raise of GFP intensity is L-DOPA concentration-dependent. Arrows indicate PPM3 clusters of TH neurons. Scale bars: (A-C) 10 μm; (E) 20 μm. (F) Quantification of GFP intensity after normalized with DsRed signals. PPM3 clusters are selected for measurement from ≥ 10 flies. Values shown represent mean ± SE. **, p < 0.01 as compared to buffer (vesicle) treated control (one-way ANOVA with Dunnett’s multiple comparisons test).

To test if the weak GFP signals result from the detection of the intrinsic cytosolic DA or other catecholamines by the probe, we co-expressed MMG1^C397A^/sfG110 in DA neurons. As compared to MMG1/sfG110 cells, GFP signals in MMG1^C397A^/sfG110 cells were undetectable (Figure 3B), supporting the notion that the weak GFP signal observed in *TH>MMG1/sfG110* was indeed responding to endogenous DA. To further confirm this, we fed *TH>MMG1/sfG110* flies with pargyline, an irreversible MAO B inhibitor that forms covalent adduct to FAD and inhibits oxidase activity (Binda et al., 2003). As compared with untreated flies, pargyline-treated flies showed very low, if any, GFP signals. Notably, increased GFP emission was observed after replacing the inhibitor with ascorbic acid buffer used to prepare pargyline and allow MMG1 protein synthesis (Figure S4). Altogether, these data suggest that the MMG1/sfG110 is feasible to detect cytosolic DA. We name this probe CyDAP for its function as a <u>cy</u>tosolic <u>DA</u> probe.

### CyDAP Reports Cytosolic DA Dynamics

To test CyDAP in responding to the cytosolic DA dynamics, we first compared the intensity of GFP emission from the probe driven by *TH-D’-GAL4,* expressed in a subset of DA neurons (Liu et al., 2012). With TH overexpression, an approach that has been shown to elevate DA production (Franco et al., 2010; Locke et al., 2008; Park et al., 2007; Vecchio et al., 2017), we found an increase in GFP intensity compared to LacZ control (Figure 3C and 3D). Because synthesized DA requires VMAT to transport into a synaptic vesicle, we tested the probe in VMAT heterozygous background (*VMAT^SH0459/+^*), a condition could hinder DA storage and increase cytosolic DA level (Mosharov et al., 2003; Mosharov et al., 2009; Vergo et al., 2007). Indeed, we found GFP emission from *TH-D’>CyDAP* neurons was enhanced in *VMAT^SH0459/+^* background as compared to control (Figure 3C and 3D), further support this probe is feasible for detecting cytosolic DA dynamics.

L-DOPA therapy is a conventional treatment in PD, as refurbishing DA could relieve the patient’s symptoms. Because a similar effect has been demonstrated in some PD fly models (Liu et al., 2008), we asked whether feeding flies with L-DOPA could increase GFP emission in *TH>CyDAP* flies. By co-expressed a reference DsRed marker with CyDAP, we found a dosage-dependent increase of GFP emission (Figure 3E and 3F). Notably, we observed that DA neurons in the so-called PPM3 cluster are most responsive to L-DOPA treatment (arrows, Figure 3E), which results coincided with the observation that PPM3 neurons are more sensitive to the insult of rotenone-treatment in a fly PD model (Coulom and Birman, 2004).

### Imaging Cytosolic DA fluctuation in Living Brains

Next, we asked whether CyDAP could detect DA fluctuation in live (detail procedure in Materials and Methods). By focusing on the somas of DA neurons in the brains of *TH-D’>CyDAP* flies, we found that GFP signals remain unchanged before and after adding the HL3 buffer that was used to dissolve the DA (Supplementary Video 3 and green lines in Figure 4B). In contrast, after adding 1 mM DA, the same DA neurons emitted GFP signals with increased intensity around ~50 folds compared to buffer-treated control (Supplementary Video 4 and Figure 4A, and red lines in 4B). On average, the signals peaked at ~3 min after DA addition and returned to baseline at ~7 min.

**Figure 4.**
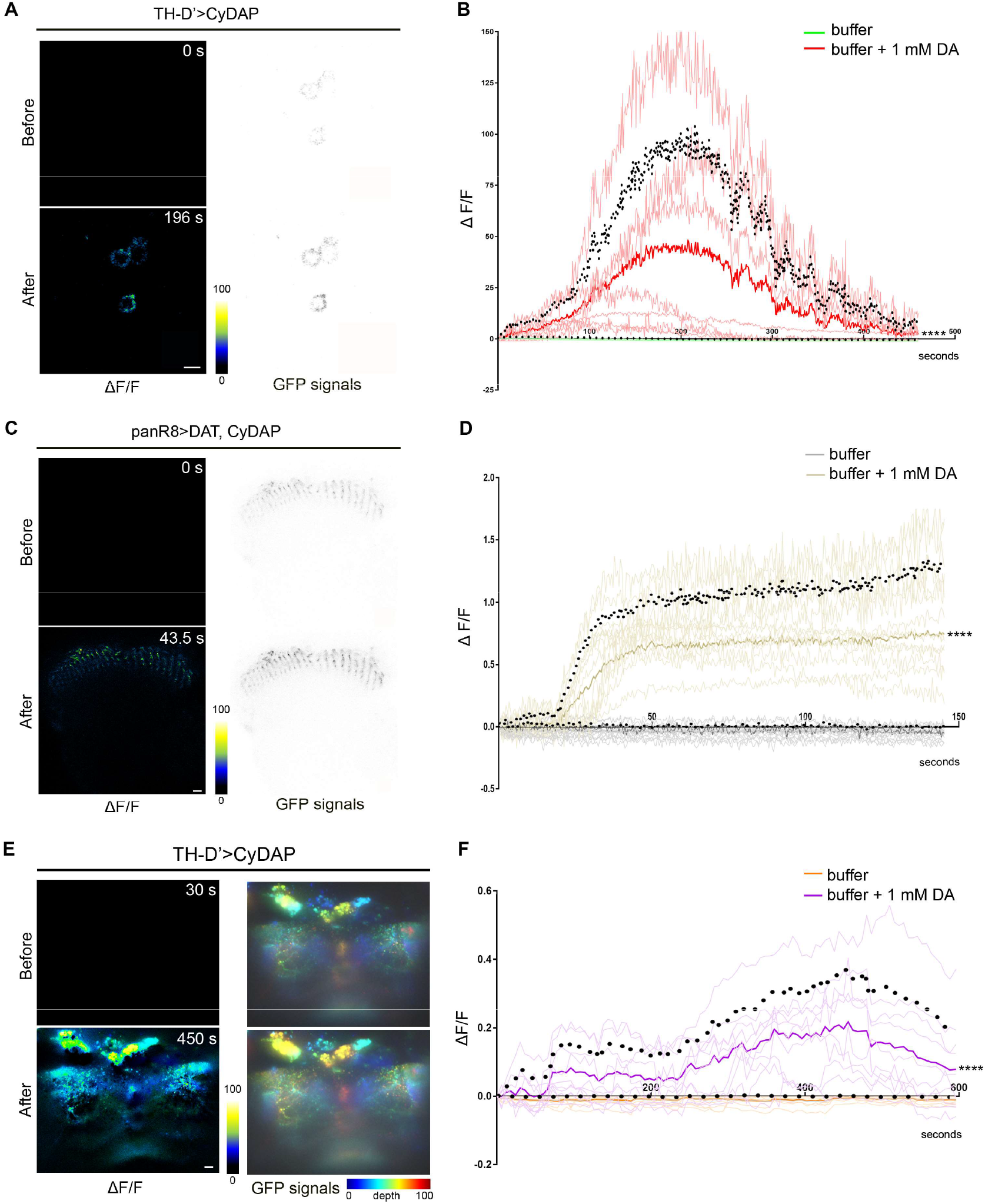
CyDAP detects DA dynamics in live imaging. (A-D) 2-D live imaging of *TH-D’>CyDAP* (A, B) and *panR8>DAT, CyDAP* (co-expressing probe and DAT; C, D), flies with or without 1 mM DA treatment. Representative images show TH-D’ neuron soma (A) and panR8 neuron axonal terminal (C) at the indicated time points, the intensity of the GFP signals is presented in grayscale for clarity (A, C, right panels). (E, F) 4D light-sheet live imaging of *TH-D’>CyDAP* flies with or without 1 mM DA treatment. Representative volumetric GFP intensities show a brain before and after DA treatment. Images are colored-coded with depth indicator processed by ImageJ-Fiji jet hyper stack in projected view (E, right panels). The ΔF/F is presented in color grading and processed by ImageJ image calculator (A, C, E, left panels); the time-lapse recordings are shown in the diagram and thick lines and black dots are mean + SD (B, D, F). The increase of GFP emission (ΔF/F) from the corresponding treatments is plotted during the recording. Brains are treated with HL3 buffer (green lines in B; n=7, gray lines in D; n=7, and orange lines in F; n=4), 1 mM DA (red lines in B; n=7, gold lines in D; n=7, and purple lines in F; n=5). Thick lines represent the mean value of each group. ****, p < 0.0001 as compared to the HL3 buffer control (one-way ANOVA with Dunnett’s multiple comparisons test). Scale bars: 10 μm.

To further validate the rise of GFP intensity was due to the uptake of the added DA, we tested the probe in cholinergic photoreceptor or GABAergic neurons by using *panR8-GAL4* (Lin et al., 2016) or *202508-GAL4* (Chi et al., 2020), respectively. By expressing CyDAP, we did not detect an evident change of GFP signals with or without DA treatment in both neuron types (Figure S5). However, by co-expressing DAT in *panR8>CyDAP,* we could observe elevated GFP emission (Supplementary Video 5 and Figure 4C, and gold lines in 4D), albeit the signals did not return to baseline during the recording. The increased GFP signals responded to DA because the probe did not show boosted GFP emission after treating GABA in GABAergic neurons or DA in non-DA neurons (Figure S5).

Finally, we analyzed the probe response to DA using the 4D light-sheet fluorescence microscope. We specifically focused on the neurites of *TH-D’-GAL4-expressed* dopaminergic neurons. By quantifying the same volume before and after DA treatment, we showed that GFP signals were increased in response to DA treatment (Supplementary Video 7 and Figure 4E, and purple lines in 4F). Altogether, these results substantiate the utility of CyDAP in detecting DA fluctuation in living tissues.

### Elevated Cytosolic DA in the Brain Expressing α-Synuclein

Expression of α-Synuclein caused selective neurotoxicity towards DA-producing neurons (Xu et al., 2002). While the *Drosophila* genome lacks α-Synuclein ortholog, expressing human α-Synuclein could cause DA neuron loss at 30-day-old flies (Feany and Bender, 2000; Mizuno et al., 2010). To this end, we set to test whether the CyDAP might detect the change when DA neurons were expressing wild-type α-Synuclein and two juvenile-onset PD mutations, A30P and A53T (Kruger et al., 1998; Mizuno et al., 2010; Polymeropoulos et al., 1997). Interestingly, we found that GFP signals of CyDAP were significantly increased in DA neurons co-expressing wild-type α-Synuclein and α-Synuclein^A30P^, but not for α-Synuclein^A53T^ despite a trend of raising, at 7-day-old flies (Figure 5). This fluorescent probe thus revealed a similar result that the expression of pathogenic or wild-type human α-Synuclein could lead to the elevation of free cytosolic DA observed by the intracellular patch electrochemistry recording (Mosharov et al., 2009; Mosharov et al., 2006). Importantly, this data also suggests that in α-Synuclein-associated PD condition, an aberrant increase of cytosolic free DA may play a pathogenic role.

**Figure 5.**
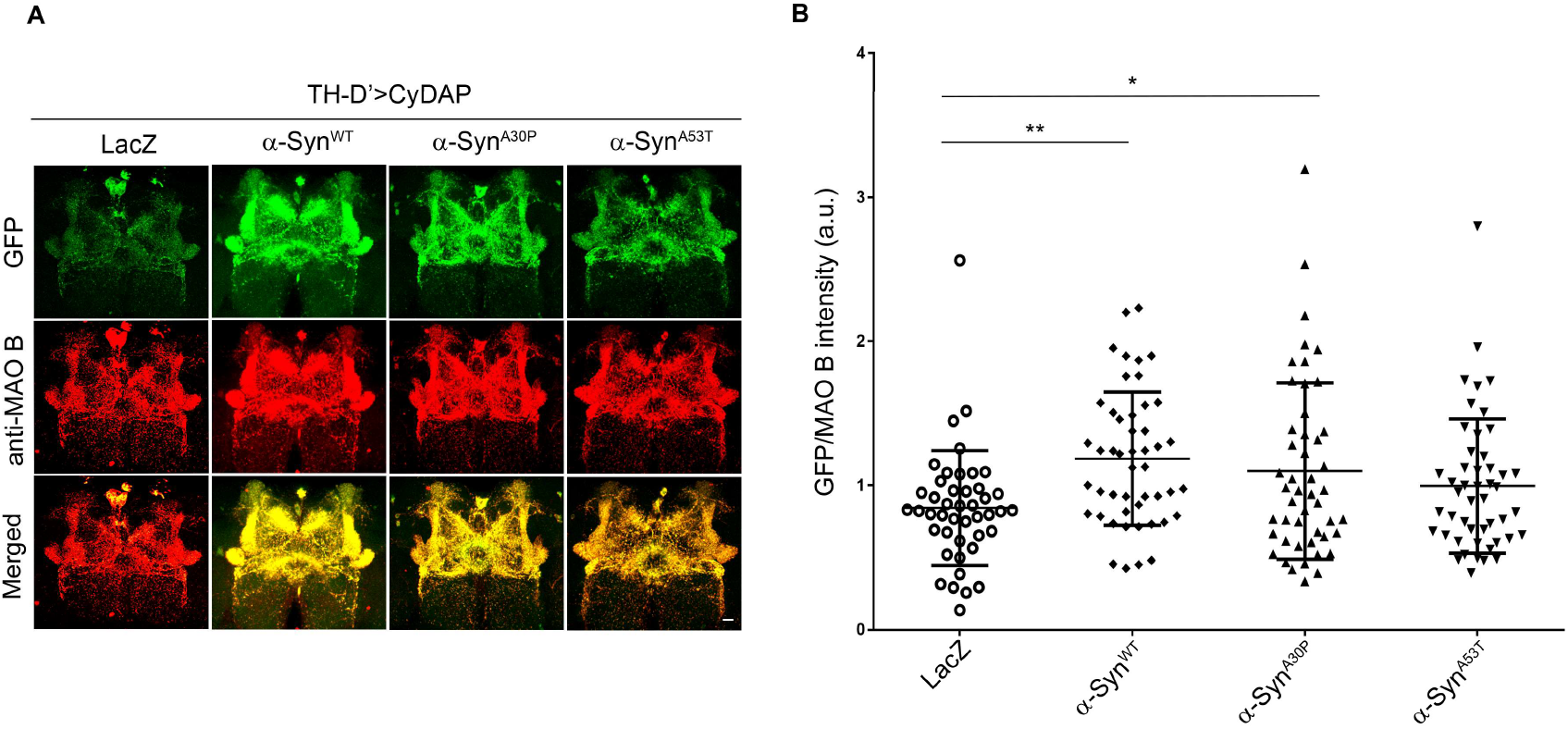
CyDAP reveals increased cytosolic DA in *Drosophila* DA neurons expressing human α-Synuclein. (A) Confocal images of the brains from *TH-D’>CyDAP* flies co-expressing LacZ control or the indicated human α-Synuclein transgenes stained with anti-MAO B (red). The intrinsic GFP signals from MMG1/sfG110 are shown in green. Scale bar: 10 μm. (B) Quantification of GFP intensity that normalized with anti-MAO B signals. Four neuropil regions are measured in each brain from a total of 12 files in each group. Values shown represent mean ± SE. *, p < 0.05; **, p < 0.01 as compared to LacZ (one-way ANOVA with Dunnett’s multiple comparisons test).

## Discussion

DA-linked cytotoxicity is an appealing mechanism in PD, particularly when we consider the disease’s regional vulnerability. However, validating this pathological prospect requires a readout of the free DA pool in affected neurons. Here, we report a fluorescent probe enabling the detection of cytoplasmic and non-compartmentalized DA. This probe, CyDAP, is a bipartite split-GFP design in which the enzymatic MMG1, modified from human MAO B by tethering an essential membrane-spanning fragment and the 11th β-sheet of GFP with limited disturbance of the enzyme activity, self-assembles with sfG110. Expressing CyDAP in cultured cells reveals the probe could detect the presence of DA. Genetic and drug manipulations of CyDAP-bearing *Drosophila* demonstrate the fluorescent responses from this probe are associated with cytoplasmic free DA dynamics, advancing its usefulness on deciphering the role of free cytosolic DA in selective nigrostriatal neuronal loss in PD.

DA is a highly reactive compound. DA synthetic enzymes, TH and DDC, forms a functional complex with VMAT at the synaptic vesicle membrane to transport the newly synthesized DA in the synaptic vesicle (Cartier-Z et al., 2010), suggesting that free cytoplasmic DA is tightly regulated. While non-compartmental DA could be metabolized into the benign homovanillic acid; indeed, free DA could metabolite into highly reactive aldehyde intermediates and generate hydrogen peroxide as by-products (Eisenhofer et al., 2004; Goldstein et al., 2013; Rees et al., 2009). Furthermore, auto-oxidation of DA under the physiological condition could form cysteinyl quinone adducts (LaVoie and Hastings, 1999; Lohr et al., 2017; Lotharius and O’Malley, 2001). Through the direct injection of DA, or via manipulating transporters functions, presumably to control cytosolic DA levels, several studies have demonstrated that those attempted alterations of cytosolic DA link to cytotoxicity (Chen et al., 2008). For instance, VMAT2-deficient in mice depleted vesicular filling of DA, which exacerbated dopaminergic neurodegeneration (Caudle et al., 2007; Fon et al., 1997; Taylor et al., 2014). Similarly, VMAT^SH0459^ loss-of-function mutant used in this study could cause dopaminergic neurons loss in *Drosophila* (Lawal et al., 2010; Simon et al., 2009). However, the link between cytosolic DA and neurotoxicity was only validated until the subcellular electrochemical probe recording (Mosharov et al., 2009). Importantly, our fluorescent probe confirms VMAT deficiency indeed increases cytosolic free DA. Furthermore, we also find that TH overexpression in DA neurons could raise cytosolic free DA levels based on the CyDAP. This data is consistent with the observation in mice because increasing TH levels by expressing multiple copies of functional transgenes could increase cysteinylated catechols and oxidative stress (Vecchio et al., 2017). It will be interesting to test whether the cytosolic free DA was the cause in this model.

α-Synuclein is the major component found in Lewy bodies or Lewy neurites, a PD cellular hallmark (Spillantini et al., 1997). Genetic variants of SNCA, the gene that encodes α-Synuclein, have been linked to familial and sporadic PD. This natively unstructured protein is enriched in the presynaptic terminals and may involve different aspects of DA homeostasis (Bridi and Hirth, 2018; Burre, 2015; Lotharius and Brundin, 2002). Previous studies found that α-Synuclein could interact with TH and DDC and thereby affecting DA biosynthesis (Perez et al., 2002; Tehranian et al., 2006). Moreover, α-Synuclein and VMAT2 could form a complex, whereas its upregulation inhibited VMAT2 activity and reduced DA uptake (Guo et al., 2008). Such regulation may instigate a harmful feedback circuit as α-Synuclein is prone to be modified by DA adducts and form protofibrils (Conway et al., 2001), ultimately resulting in oxidative stress and neurodegeneration (Lotharius and Brundin, 2002; Park et al., 2007). Our probe detects increased cytosolic free DA upon expressing human α-Synuclein, consistent with the finding using the electrochemical probe (Mosharov et al., 2009).

Overall, we have demonstrated that our probe can detect DA dynamic in *Drosophila,* but further investigation is required for its application in mammals. Unlike flies, mammals express MAO B; thus, the cellular impact of this probe requires some pilot tests. Reports in mouse and human showed that MAO B level is associated with Alzheimer’s disease (Schedin-Weiss et al., 2017), anxiety (Schalling et al., 1987), and oxidative damage in neurons and hearts (Kaludercic et al., 2014; Kumar et al., 2003). With an appropriate experimental design, we anticipate that CyDAP could be applied to other genetic models to expand the investigation on the impacts of PD risk factors in DA neuron vulnerability.

## Materials and Methods

### DNA constructs

Human MAO B cDNA was obtained from Invitrogen (IOH55406), AcGFP was obtained from Clontech (632489), and mitofusin 2 (Mfn) was obtained from OriGene (SC114726). To generate MG constructs, MAO B of corresponded cDNA fragments were amplified by PCR to generate SacI and AgeI sites and introduced into pET23a. AcGFP was subcloned into the C-terminal of truncated MAO B as AgeI-NotI fragment and the MAO-GFP fragment was subcloned into pPyCAGIP (Chambers et al., 2003) with XhoI and NotI sites. For MG-m and MG-s, AcGFP^7-238^ was amplified by PCR to substitute full-length AcGFP used in MG-l instead.

For MMG1, full-length MAO B was PCR amplified as an XhoI-AgeI fragment. The second transmembrane domain of Mfn and split GFP11 were amplified by PCR and subsequently cloned into pPyCAGIP. MMG1^C397A^ mutation was introduced into MMG1 using the QuikChange Lightning Site-Directed Mutagenesis kit (Agilent). For MMGFP, full-length AcGFP was amplified and introduced into MMG1 to replace the GFP11 using BglII and NotI sites. As for sfGM, split GFP11 was amplified and then subcloned into the N-terminal of MAO B with XhoI and NheI sites. Split GFP110 fragment was PCR amplified and introduced into pPyCAGIP with XhoI and NotI sites. Primers used in the abovementioned constructs are listed in Supplementary Table 1. All constructed were verified by DNA sequencing.

### Cell culture

PC12 cells were plated on collagen I (BD) coating petri dishes and cultured in DMEM medium (Gibco), supplemented with 10% horse serum, 5% fetal bovine serum, and 1X antibiotics (Gibco). As for HEK293 and HEK293T, cells were cultured in DMEM supplied by 10% fetal bovine serum and 1X antibiotics on uncoated petri dishes. To differentiate PC12 cells, mediums containing 1% horse serum, 0.5% fetal bovine serum, 1X antibiotics, and 80ng β-NGF/ml (BD) were used. Transfection was carried out in culture dish with 80% confluence by using Lipofectamine 2000 (Invitrogen) or jetPRIME (Polyplus) following the manufactures’ instructions.

### Western blotting

Protein lysates were collected from cells and extracted with lysis buffer (20 mM Tris, 150 mM NaCl, 1 mM EDTA, 1% Triton X-100, 2.5 mM sodium pyrophosphate, 1 mM β-glycerophosphate and 1X protease inhibitor cocktail (Roche). Proteins extracts were resolved on 4-12% Bis-Tris NuPAGE (Invitrogen). Primary antibodies used were rabbit anti-GFP 1:2000 (Invitrogen), rat anti-DAT 1:2000 (Chemicon), and mouse anti-actin 1:10000 (Novus biological). Secondary antibodies conjugated with HRP (Jackson ImmunoResearch Laboratories) were used in 1:10,000 dilutions. All loading controls were prepared by stripping off the reagents from the original membrane and then re-probed by following the standard procedures.

### Flow cytometry

The transfected PC12 cells were incubated with or without benzylamine at 37°C incubator for 30 min. Cells were harvested and resuspended with 5 ml DMEM. After PBS washes, cells were resuspended in 2% paraformaldehyde and incubated on ice for 20 min. The fixed cells were washed and resuspended in PBS with a final concentration of 0.5×10^6^ cells/ml. GFP-positive cells were counted using BD Accuri C6 flow cytometry.

### Cell injection

Cells were plated on 35 mm coated petri dishes and cultured in medium containing 20 ng/ml NGF for 4 days. The culture medium was replaced by KRH buffer (25 mM HEPES, 125 mM NaCl, 4.8 mM KCl, 1.2 mM KH_2_PO_4_, 1.3 mM CaCl_2_, 1.2 mM MgSO_4_, 5.5 mM glucose, pH 7.4) before the injection. Fresh prepared DA was delivered by Borosilicate micropipette (O.D.=1.0 mm, I.D.=0.5 mm) under the control of a Pneumatic PicoPump (World Precision Instrument). A CCD camera equipped in ZEISS axioskop2 FS+ inverted microscope with a water immersion lens recorded the videos.

### Drug treatments

For L-DOPA and BZA treatments, 3,4-Dihydroxy-L-phenylalanine and Benzylamine (Sigma) of the indicated concentrations were freshly prepared in KRH buffer containing 1 mM ascorbic acid as the anti-oxidant before the experiment. Cells subject for the treatment were incubated for 20 min before 1X PBS washing and fixation. Flies subject for the treatment were fed with yeast-glucose-agar medium (8% yeast, 8% glucose, and 1.6% agar) containing the indicated final L-DOPA concentrations. The treatment of Pargyline in flies followed the same preparation where the indicated final concentration of pargyline hydrochloride was dissolved in the fly medium. For controls, samples were treated with KRH buffer containing ascorbic acid.

### Immunohistochemistry

For cultured cells, cells were seeded on Matrigel (BD)-coated coverslips. Cells were rinsed with 1X PBS before the fixation with 4% paraformaldehyde (Sigma) for 10 min, followed by permeabilization with PBST (0.1% Triton X-100). For flies, the brains were dissected and fixed in 4% paraformaldehyde (Sigma) for 30 min, followed by permeabilization with PBST (0.3% Triton X-100). The primary antibodies used were rabbit anti-MAO B (1:200, GeneTex), goat anti-GFP (1:500, Abcam), mouse anti-ATP synthases (1:500, Molecular Probes), mouse anti-6xHis (1:600, GeneTex), mouse anti-ATP5a (1:500, Abcam), mouse anti-TH (1:200, Immunostar), and rat anti-DAT (1:200, Millipore). To label mitochondria in culture cells, MitoTracker Red CXRos (Molecular Probes) was used in 1:2500 dilution. Alexa Fluor 488, Cy3, and Cy5 conjugated secondary antibodies (Jackson ImmunoResearch Laboratories) were used at 1:1000 (cells) or 1:400 (brains) dilutions. Samples were mounted with antifade mounting medium (Vectashield H-1000, Vector Laboratories). All fluorescent images were collected on a ZEISS LSM510 confocal microscope under the same imaging setting for each set of experiments. Images were processed with Photoshop 2020.

### MAO B activity assay

MAO B catalytic activity was determined by MAO-Glo Assay (Promega). Briefly, HEK293 cells were seeded on 96-well culture plate and transfected with the indicated plasmids for 24 hrs. The reaction of cultured cells and MAO substrate were performed in MAO B reaction buffer (100 mM HEPES, pH 7.5, 5% glycerol, and 10% dimethyl sulfoxide) at 37°C for 30 min. Afterward, 50 μl of reconstituted Luciferin Detection Reagent was added for each reaction, and the mixtures were incubated at room temperature for 20 min to terminate the reaction. MAO B oxidized the amino group of luminogenic MAO substrate to produce methyl luciferin, which was next converted into light by the esterase and luciferase. The luminescent intensity was measured with VICTOR 3 (PerkinElmer), and the values were normalized to the MAO B protein level measured by Western blotting.

### Mitochondria Isolation and MAO B Kinetics

Approximately 2×10^7^ cells were harvested by centrifugation (600 g, 10 min at 4°C). After PBS wash with and re-centrifugation, the pellet was resuspended in 400 μl Solution A (250 mM sucrose, 0.1 M KH_2_PO_4_, 1x protease inhibitor, pH 7.4) and homogenized with a loose glass-glass tissue grinder. The homogenate was subsequently centrifuged for 10 min at 800 g to remove the cell debris and nuclei. The supernatant was centrifuged (16,000 g, 20 min at 4°C) to yield a pellet of crude mitochondria fraction. The pellet was then resuspended with 100 μl Solution A and treated with BZA to calculate substrate affinity using a spectrophotometric assay.

### *Drosophila* genetics

Flies were maintained at 25°C on standard cornmeal media in 12 hr light/dark cycles. *TH-D’-GAL4* was a gift from Dr. Mark Wu (Johns Hopkins University), *panR8-GAL4* was provided by Dr. Chi-Hon Lee (Academia Sinica). *TH-GAL4, UAS-DAT, VMAT^SH0459^,* and human α-Synuclein flies were obtained from the Bloomington *Drosophila* stock center. *202508-Gal4* was obtained from the Vienna *Drosophila* Resource Center. *UAS-TH* was a gift from Dr. Tsai-Feng Fu (National Chi Nan University). To generate transgenic flies, the abovementioned pPyCAGIP plasmids of MMG1, MMG1^C397A^, sfGM, and sfG110 were subcloned into the *Drosophila* transformation vector pUAST using the EcoRI-NotI fragments. All constructs were sequencing verified before transgenic fly production.

### Live imaging

Fresh dissected fly brains were adhered to slides by worm glue (GluStitch) before soaked in HL3 buffer (70 mM NaCl, 5 mM KCl, 20 mM MgCl_2_, 10 mM NaHCO_3_, 5 mM trehalose, 115 mM sucrose, 5 mM HEPES, pH 7.3). 40x water immersion lens (ZEISS, 1.2 NA, 421767-9971-711) was used to imaging the tissues. DA was freshly prepared by dissolving dopamine hydrochloride (Abcam) in HL3 buffer with 1 mM ascorbic acid. DA was gently added in the HL3 buffer at the final concentration of 1 mM during the image acquisition. For the brain, we acquired images in a speed of 75 frames/min (TH-D’ group) and 120 frames/min (R8 and 202508 groups). The time-lapse recordings were assembled into video clips via the onboard Zen software of ZEISS LSM-780 confocal microscope. The GFP intensity was determined by ImageJ and plotted by using Prism.

For 4D light-sheet imaging, individual flies with an open on the head cuticle that exposes a part of the brain that was sticking to a steel plate and soaking in the observing chamber filled with HL3. The image was taken with customized light-sheet microscopy. The excitation laser is 488 nm (Coherent OBIS 488 nm LS 150 mW). The laser beam used an acousto-optic tunable filter (AOTF; AOTFnC-400.650-TN, AA Quanta Tech, Optoelectronic) to control the exposure time and wavelength selection. The Gaussian intensity distribution of the laser was projecting to an annular ring pattern on the customized aluminum coating mask (thickness of 1,500 angstroms). The masked laser ring image was then projected to a set of galvanometer scanners (6215H, Cambridge Technology), which are composed of a pair of achromatic lenses (Thorlabs, AC254-100-A, Achromat, Ø1”, 400–750 nm), aligned in a 4f arrangement. After passing through the scanning mirror set, the ring pattern is magnified through a relay lens (Thorlabs, AC254-250-A and AC254-350-A Ø1” Achromat, 400–750 nm) and conjugated to the back focal plane of the excitation objective (Nikon CFI Plan Fluorite Objective, 0.30 NA, 3.5 mm WD). The annular pattern was projected to the rear focal plane of the excitation objective and forms a self-reconstructive Bessel beam by optical interference. We used a water dipping objective lens (Nikon, CFI Apo LWD 25XW, 1.1 NA, 2 mm WD for imaging in PBS) as the detection objective, which orthogonal to the illumination plane. We mounted on a piezo scanner (Physik Instrumente, P-725.4CD PIFOC), used to collect the fluorescence signal, which then passes an emission filter (Semrock Filter: FF01-446/523/600/677-25) onto a complementary scientific metal-oxide-semiconductor (sCMOS) camera (Hamamatsu, Orca Flash 4.0 v3 sCOMS) by a 250 mm tube lens. We imaged 30 frames through the SDFP region of the fly brain for each time point, and each frame was taking in 30 ms, 300 um x 300 um. Imaris 9.1.1. (Bitplane AG) was used to generate 4D movies of how DA neurons respond to DA treatment. We converted the 3D tiff files for each time point into 4D time-series data in ims format with Imaris file converter. We then used the Animation function to generate 4D movies.

## Acknowledgements

This study was supported by grants from NTHU-NTUH joint project (103N2778E1 and 104N2276E1) and the Higher Education Sprout Project funded by the Ministry of Science and Technology and the Ministry of Education in Taiwan. We thank Drs. Chi-Hon Lee, Mark Wu, Tsai-Feng Fu, Bloomington *Drosophila* Stock Center, Vienna *Drosophila* Resource Center and Fly Core in Taiwan for generously providing fly strains. We thank Dr. Chuan-Chin Chiao for live-cell imaging and the Image Core of the Brain Research Center at National Tsing Hua University for assistance with confocal and 4D light-sheet microscopy.

