## supplementary figures for "CyDAP–A fluorescent probe for cytosolic dopamine detection"

### Supplementary Figure Legends

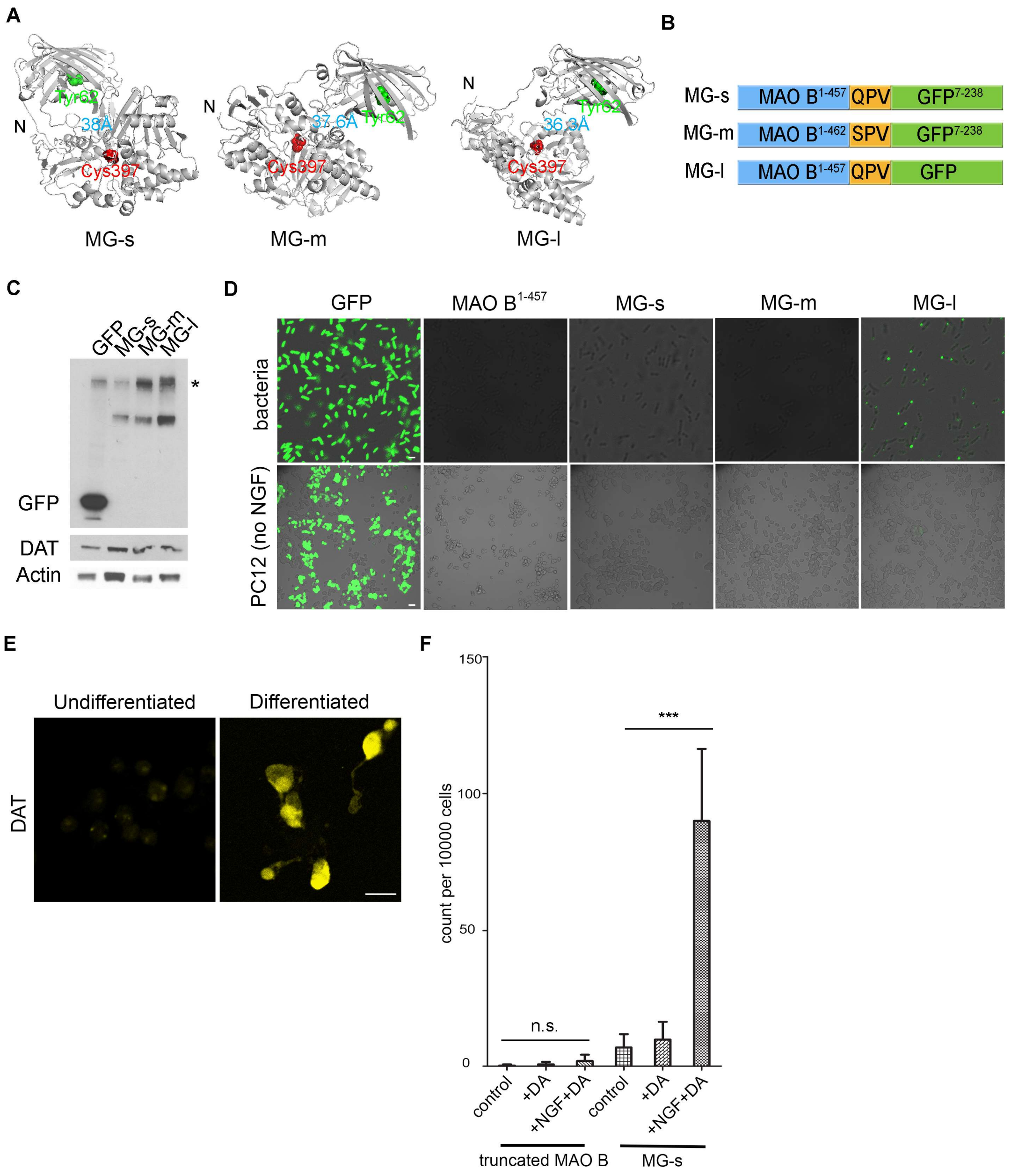


Figure S1

**Supplementary Figure 1. The constructs of the tested DA probes.** (A) The prediction of MG-s, MG-m, and MG-l protein structures (https://zhanglab.ccmb.med.umich.edu/I-TASSER/). PyMOL labels the distance (blue) between the FAD-binding residue Cys397 of MAO B (red in ball-and-stick) and one of the fluorophore-composing residues of GFP, Tyr62 (green in ball-and-stick). (B) Schematic depiction of the designed constructs with colored blocks to mark the corresponded MAO B (blue), linker residues (yellow), and GFP (green). (C) Western analysis of protein extracts from differentiated PC12 cells that are transfected by the indicated constructs. The full-length GFP serves as a control. The detected GFP, DAT, and actin are marked. The asteroid on the top blot indicates a non-specific band. (D) Fluorescent images merged with bright field show *E. coli* bacteria and undifferentiated PC12 cells that are transfected by the indicated constructs. (E) Confocal images of PC12 cells before (undifferentiated) and after treating with NGF (differentiated) that are labeled by the anti-DAT antibody. (F) Quantitative analysis of GFP-positive PC12 cells transfected with truncated MAO B and MG-s and the indicated treatments. Values shown represent mean ± SE. n.s. not significant; ***, $p < 0.001$ as compared to control (n=3, Student's t-test). Scale bars: (D) 20 μm, (E) 10 μm.

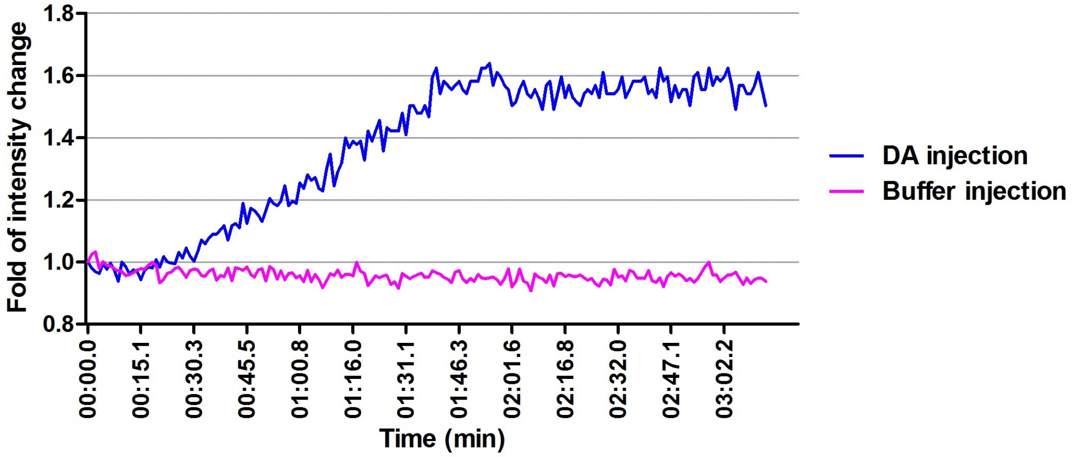


Figure S2

**Supplementary Figure 2. Recording of live PC12 cells transfected by MG-s with DA injection.** The time-course plot of GFP intensity from PC12 cells transfected by MG-s are injected with KRH buffer (magenta line) or DA (blue line) shown in Supplementary Video 1 and 2).

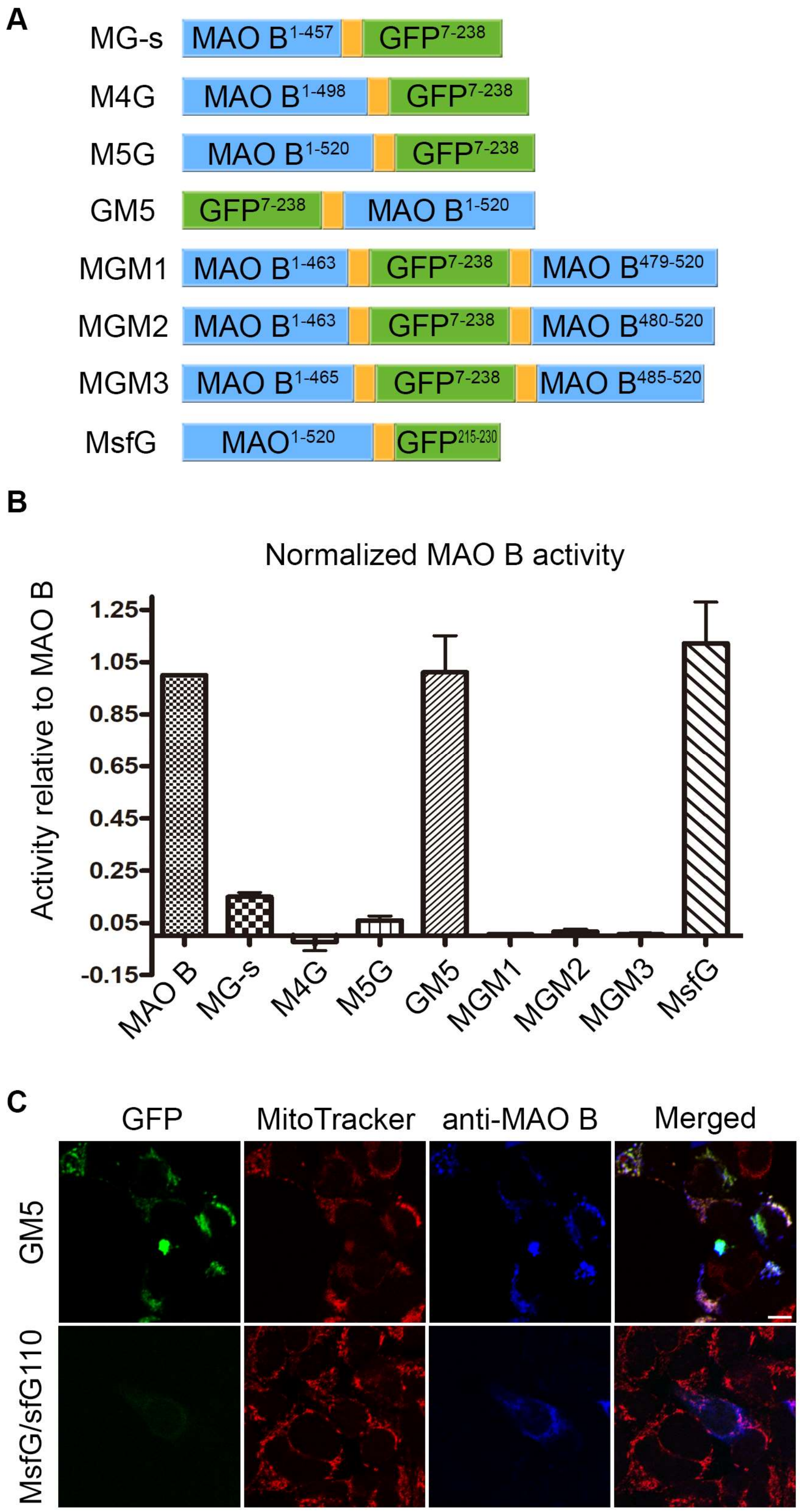
A
MG-s
MAO B1-457
GFP7-238
M4G
MAO B1-498
GFP7-238
M5G
MAO B1-520
GFP7-238
GM5
GFP7-238
MAO B1-520
MGM1
MAO B1-463
GFP7-238
MAO B479-520
MGM2
MAO B1-463
GFP7-238
MAO B480-520
MGM3
MAO B1-465
GFP7-238
MAO B485-520
MsfG
MAO1-520
GFP215-230
B
Normalized MAO B activity
Activity relative to MAO B
1.25
1.05
0.85
0.65
0.45
0.25
0.05
-0.15
MAO B
MG-s
M4G
M5G
GM5
MGM1
MGM2
MGM3
MsfG
C
GFP
MitoTracker
anti-MAO B
Merged
GM5
MsfG/sfG110

Figure S3

**Supplementary Figure 3. DA probe modifications.** (A) Schematic diagrams of the tested DA probe constructs. The corresponded MAO B (blue block) and GFP (green block) segments are indicated. Yellow blocks denote linker regions. (B) Quantitative analysis of MAO B activity of the indicated constructs. Values shown represent mean ± SE as compared to full-length MAO B (n=3). (C) Confocal images of HEK293 cells transfected with GM5 or MsfG/sfG110 stained with MitoTracker (red) and anti-MAO B (blue). Notice GFP signals in GM5, but not in MsfG/sfG110. Scale bar: 10 μm.

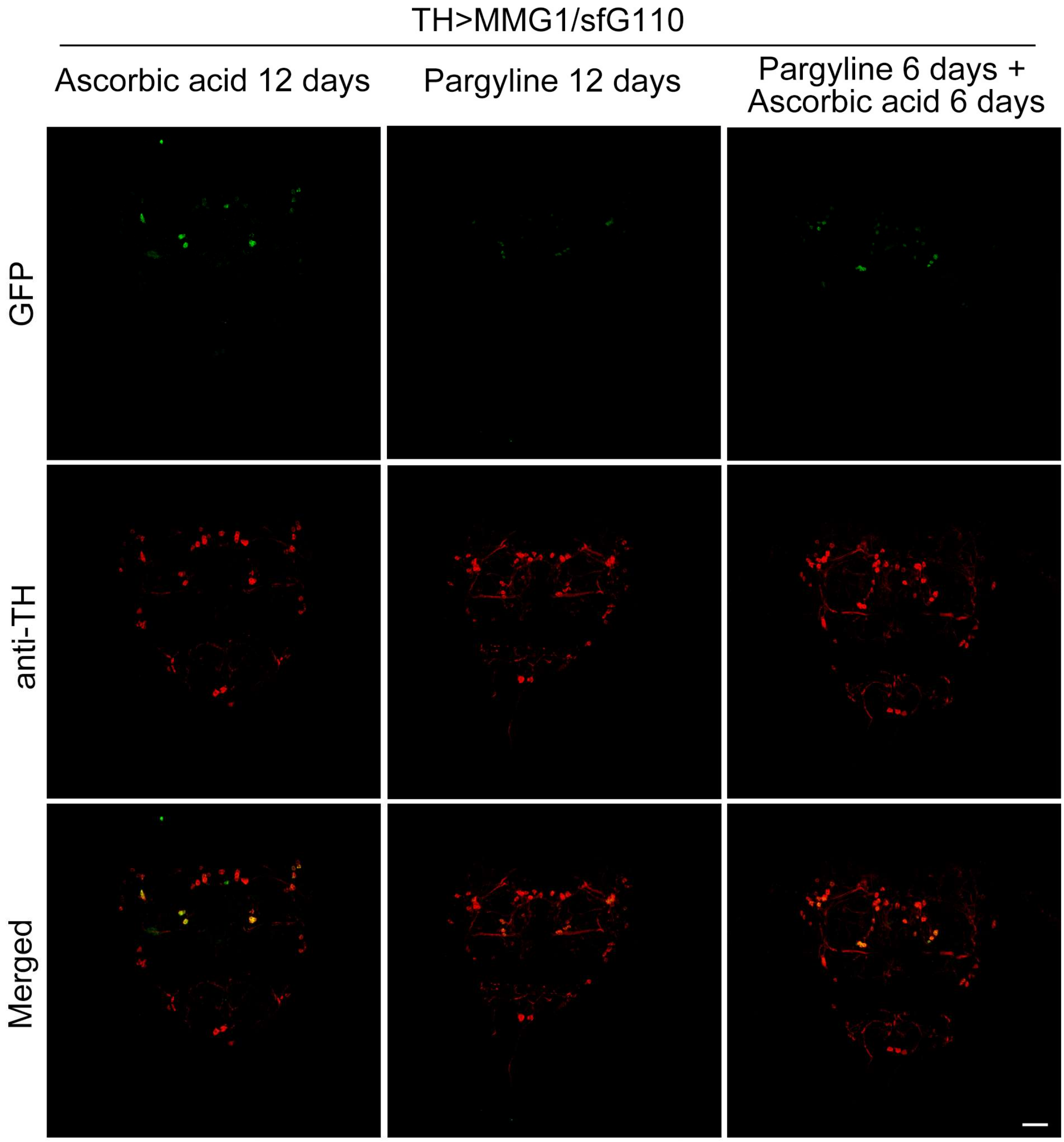


Figure S4

**Supplementary Figure 4. The treatment of the MAO B inhibitor blocks GFP emission in DA neurons expressing MMG1/sfG110**. Confocal images of the brains from *TH>MMG1/sfG110* flies treated with the indicated remedies stained with anti-TH (red). The GFP signals are intrinsic from the probe. Ascorbic acid is an antioxidant used in the buffer that dissolving the MAO B inhibitor pargyline. Scale bar: 20 μm.

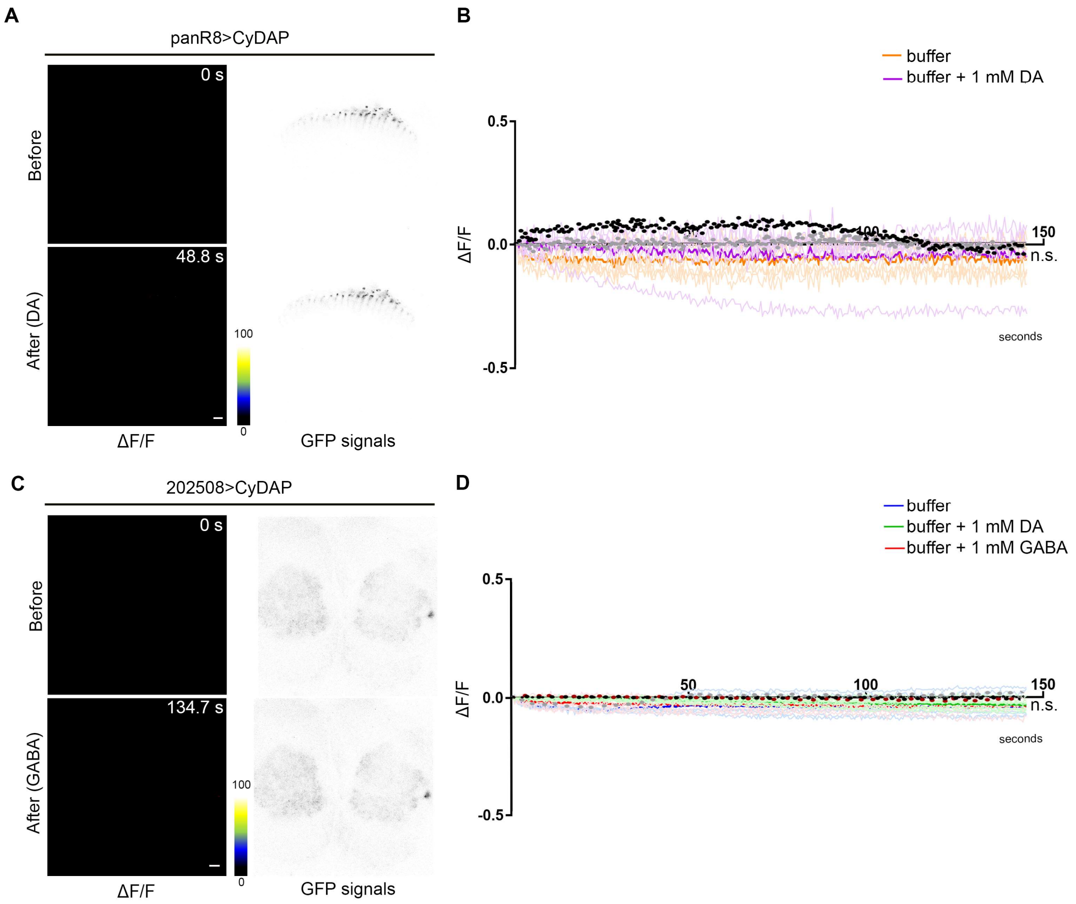


Figure S5

**Supplementary Figure 5. Live imaging of non-DA neurons expressing the CyDAP.** (A-D) 2-D live imaging of *panR8>CyDAP* (A, B) and *202508>CyDAP* (C, D) before and after 1 mM DA or 1 mM GABA treatments. Representative images show R8 neuron axonal terminal (A) and antennal local neuron axonal terminal (C) at the indicated time points. The GFP signal intensity is presented in grayscale for clarity (A, C, right panels). The ΔF/F is presented in color grading (A, C, left panels) and processed by ImageJ image calculator. The time-lapse recordings are shown in the diagram and thick lines and black dots are mean + SD (B, D). Brains are treated with HL3 buffer (orange lines in B; n=3, blue lines in D; n=4), 1 mM DA (purple lines in B; n=3, green lines in D; n=4), and 1 mM GABA (red lines in D; n=3). There is no significant difference compared to control (one-way ANOVA, Dunnett's multiple comparisons test). Scale bars: 10 μm.

### Supplementary Video Legends

**Supplementary Video 1 & 2. Recording of live PC12 cells transfected by MG-s with DA injection.** The GFP signals from PC12 cells transfected by MG-s are recorded by ZEISS axioskop2 $FS^+$ inverted microscope with a water immersion lens. Cells are injected with 100 μM DA (Video 1) or KRH buffer as control (Video 2). The videos are in six-time speed and edit by Power Director 15.

**Supplementary Video 3 & 4. CyDAP responses to DA in *Drosophila* dopaminergic neurons in the brain.** Live imaging of *TH-D'>CyDAP* flies with (Video 4) or without (Video 3) 1 mM DA treatment. Images are acquired by ZEISS LSM-780 confocal microscope with a 40x water immersion lens. Videos are exported by ZEN in 20 frames/second and edit by Power Director 15.

**Supplementary Video 5 & 6. CyDAP responses to DA in *Drosophila* cholinergic neurons.** Live imaging of optic lobes from *panR8>DAT, CyDAP* (co-expressing probe and DAT) flies with (Video 5) or without (Video 6) 1 mM DA treatment. Images are acquired by ZEISS LSM-780 confocal microscope with a 40x water immersion lens. Videos are exported by ZEN in 20 frames/second and edit by Power Director 15.

**Supplementary Video 7 & 8. CyDAP responses to DA in volumetric projection view of the fly brain.** 4D light-sheet live imaging of *TH-D'>CyDAP* flies with (Video 7) or without (Video 8) 1 mM DA treatment. Images are acquired by customized light-sheet microscopy with a 40x water immersion lens (Nikon). Videos are converted by Imaris in 60 frames/second and editing by Power Director 15.
