## supplementary table for "CyDAP–A fluorescent probe for cytosolic dopamine detection"

**Supplementary Table 1**

| Construct | Primer | Sequence (5'-3') |
|---|---|---|
| MG-s | MAO1-SacI-F | GTACAAAAAAGAGCTCACCATGAGCAAC |
| | MAO457-AgeI-R | GCAGAATTACCGGTTGAATCTTCC |
| | GFP7-Age-F | CAACCGGTCGAGCTGTTCAC |
| | GFP238-NotI-R | GCGGCCGCTTAGTACAGCTCATC |
| MG-m | MAO1-SacI-F | GTACAAAAAAGAGCTCACCATGAGCAAC |
| | MAO462-AgeI-R | TGACCGGTTGAATCTTCCCCAT |
| | GFP7-Age-F | CAACCGGTCGAGCTGTTCAC |
| | GFP238-NotI-R | GCGGCCGCTTAGTACAGCTCATC |
| MG-l | MAO1-SacI-F | GTACAAAAAAGAGCTCACCATGAGCAAC |
| | MAO457-AgeI-R | GCAGAATTACCGGTTGAATCTTCC |
| | GFP1-AgeI-F | ACCGGTCATGGTGAGCAAGGGCG |
| | GFP238-NotI-R | GCGGCCGCTTAGTACAGCTCATC |
| M4G | MAO1-XhoI-F | TTTCTCGAGATGAGCAACAAATGCGACGT |
| | MAO498-AgeI-R | TTTACCGGTTTACAATCCAATCAGCCTGAGC |
| | GFP7-Age-F | CAACCGGTCGAGCTGTTCAC |
| | GFP238-NotI-R | GCGGCCGCTTAGTACAGCTCATC |
| M5G | MAO1-XhoI-F | TTTCTCGAGATGAGCAACAAATGCGACGT |
| | MAO520-AgeI-R | TTTACCGGTTTAGACTCTCACAAGTAGCCCCC |
| | GFP7-Age-F | CAACCGGTCGAGCTGTTCAC |
| | GFP238-NotI-R | GCGGCCGCTTAGTACAGCTCATC |
| GM5 | GFP7-XhoI-F | TTTCTCGAGGGTATGGAGCTGTT |
| | GFP238-NheI-R | TTTGCTAGCTCACTTGTACAG |
| | MAO1-NheI-F | TTTGCTAGCATGAGCAACAAATGC |
| | MAO520-NotI-R | TTTGCGGCCGCTTAGACTCTCACAAGTAGCCCC |
| MGM1 | MAO1-XhoI-F | TTTCTCGAGATGAGCAACAAATGCGACGT |
| | MAO463-AgeI-R | TTTACCGGTTGCCAGATTTCATCCTCTGGAATCTTCC |
| | GFP7-Age-F | CAACCGGTCGAGCTGTTCAC |
| | GFP238-NheI-R | TTTGCTAGCTCACTTGTACAG |
| | MAO479-NheI-F | TTTGCTAGCCTCGAGACCACCTTTTTGGAGAGACATTTG |
| | MAO520-NotI-R | TTTGCGGCCGCTTAGACTCTCACAAGTAGCCCC |
| MGM2 | MAO1-XhoI-F | TTTCTCGAGATGAGCAACAAATGCGACGT |
| | MAO463-AgeI-R | TTTACCGGTTGCCAGATTTCATCCTCTGGAATCTTCC |
| | GFP7-Age-F | CAACCGGTCGAGCTGTTCAC |
| | GFP238-NheI-R | TTTGCTAGCTCACTTGTACAG |
| | MAO480-NheI-F | TTTGCTAGCCTCGAGACCTTTTTGGAGAGACATTTGCCC |
| | MAO520-NotI-R | TTTGCGGCCGCTTAGACTCTCACAAGTAGCCCC |
| MGM3 | MAO1-XhoI-F | TTTCTCGAGATGAGCAACAAATGCGACGT |
| | MAO465-AgeI-R | TTTACCGGTTGTGACTGCCAGATTTCATCCTCTG |
| | GFP7-Age-F | CAACCGGTCGAGCTGTTCAC |
| | GFP238-NheI-R | TTTGCTAGCTCACTTGTACAG |
| | MAO485-NheI-F | TTTGCTAGCCTCGAGCATTTGCCCTCCGTGCCA |
| | MAO520-NotI-R | TTTGCGGCCGCTTAGACTCTCACAAGTAGCCCC |
| MsfG | MAO1-XhoI-F | TTTCTCGAGGCTAGCATGAGCAACAAATGCGACGT |
| | MAO520-AgeI-R | TTTACCGGTTTAGACTCTCACAAGTAGCCCCC |
| | sfG11-AgeI-F | CCCACCGGTAAGCGCGATCACATGATCTAC |
| | sfG11-NotI-R | TTTGCGGCCGCACCCCTCAGATGGCG |
| sfGM | sfG11-XhoI-F | TTTCTCGAGATGCGTGACCACATGGTCCTTCATGAG |
| | sfG11-NheI-R | TTTGCTAGCTGTAATCCCAGCAGCATTTACATACTC |
| | MAO1-NheI-F | TTTGCTAGCATGAGCAACAAATGC |
| | MAO520-NotI-R | TTTGCGGCCGCTTAGACTCTCACAAGTAGCCCC |

| | | |
|---|---|---|
| MMG1 | MAO1-XhoI-F | TTTCTCGAGGCTAGCATGAGCAACAAATGCGACGT |
| | MAO520-AgeI-R | TTTACCGGTTTAGACTCTCACAAGTAGCCCCC |
| | Mfn627-AgeI-F | TTTCTCGAGACCGGTAAGGCAGTGGGCTGGC |
| | Mfn648-Bgl2-R | TTTGCGGCCGCAGATCTCTCATAGACGTAGAGGAGGCC |
| | sfG11-Bgl2-F | CCCAGATCTCGTGACCACATGGTCCTTCATGAG |
| | sfG11-NotI-R | TTTGCGGCCGCTTATGTAATCCCAGCAGCATTTACATAC |
| MMG1H | MAO1-XhoI-F | TTTCTCGAGGCTAGCATGAGCAACAAATGCGACGT |
| | MAO520-AgeI-R | TTTACCGGTTTAGACTCTCACAAGTAGCCCCC |
| | Mfn627-AgeI-F | TTTCTCGAGACCGGTAAGGCAGTGGGCTGGC |
| | Mfn648-Bgl2-R | TTTGCGGCCGCAGATCTCTCATAGACGTAGAGGAGGCC |
| | sfG11-Bgl2-F | CCCAGATCTCGTGACCACATGGTCCTTCATGAG |
| | sfG11-6xHis-R | CACCACCACCACCACCACTGTAATCCCAGCAGC |
| | sfG11-NotI-R2 | TTTGCGGCCGCTTACACCACCACCACCACCACTGTAAT |
| MMG1$^{C397A}$ | MMG1$^{C397A}$-F | CTCTGGGGGCGCCTACACAACTTATTTCCCCC |
| | MMG1$^{C397A}$-R | GGGGGAAATAAGTTGTGTAGGCGCCCCCAGAG |
| sfG110 | sfG110-XhoI-F | TTTCTCGAGATGTCCAAAGGAGAAGAACTGTTTACC |
| | sfG110-NotI-R | TTTGCGGCCGCTTATGTTCCTTTTTCATTTGGATCTTTGC |
